# Human Cerebral Cortex Organization Characterized by Functional PET-FDG “Metabolic Connectivity”

**DOI:** 10.64898/2026.02.15.706044

**Authors:** Penghui Du, Sean E. Coursey, Ting Xu, Sharna Jamadar, Sara Nolin, Bin Wan, Hsiao-Ying Wey, Jonathan R. Polimeni, Julie C. Price, Quanying Liu, Jingyuan E. Chen

## Abstract

**Purpose:** In this study, we characterize the spatiotemporal organization of resting-state metabolic connectivity (RSMC) in the human brain, as measured by [^18^F]- fluorodeoxyglucose (FDG) functional PET (fPET-FDG). We examine the relationship between RSMC organization and resting-state functional connectivity (RSFC) derived from functional magnetic resonance imaging and other known cortical organizational principles.

**Methods:** Resting-state fPET-FDG data from 24 individuals were obtained from a publicly available repository. We characterized local metabolic organization using connectivity-based boundary mapping, with adaptations to account for the low signal-to-noise ratio of fPET-FDG data. We then estimated global metabolic organization through community detection-based network and principal gradient analyses. Furthermore, we examined how metabolic connectivity is shaped by temporal-frequency-specific components of fPET-FDG signal. Finally, we contextualized metabolic organization by relating metabolic gradients to anatomical, functional, and energetic reference measures.

**Results:** At the local scale, boundary mapping results indicated structured transitions shaped by a combination of both fast and slow fPET-FDG signals, partly overlapping with RSFC boundary maps. Globally, RSMC analyses revealed a robust metabolic structure organized along a superior-inferior cortical gradient. This pattern remained consistent across network community detection and principal gradient analyses and was primarily driven by low-frequency, minute-scale fPET-FDG dynamics. The identified large-scale metabolic profile aligns closely with several known anatomical and energetic constraints.

**Conclusion:** This study characterizes the spatiotemporal organizational principles of RSMC, deepening insight into the brain’s energetic framework and providing a basis for future cognitive and clinical investigations of metabolic connectivity organization.

## Introduction

The human brain exhibits a complex functional organization, reflected in both its continuous topographic organization and distributed inter-regional relationships. Mapping this large-scale cortical organization has been a central objective in human neuroimaging during the past decades with functional magnetic resonance imaging (fMRI) playing a dominant role through blood-oxygenation-level-dependent (BOLD) contrast [1–3]. The advantages of BOLD-fMRI includes its non-invasive nature, whole-brain coverage, milimeter-scale spatial resolution, and temporal resolution typically on the order of seconds [4,5]. In particular, resting-state functional connectivity (RSFC), derived from the temporal synchrony of BOLD-fMRI signals across distributed brain regions in the absence of an explicit task, has provided a powerful framework for characterizing large-scale brain organization [6–9]. Through RSFC, recurring patterns of inter-regional coordination have been identified, which align with known anatomical and neurochemical pathways [8,9] and task-evoked activation patterns [10–16]. To further examine principles of cortical organization across global and local scales, a range of analytical frameworks have been developed, including approaches that delineate local areal boundaries [17–20], characterize distributed functional networks [21–24], and capture continuous gradients of functional variation [25].

Despite its widespread use in human brain mapping, brain function is supported by multiple interacting physiological processes. BOLD-fMRI alone is insufficient to capture its full complexity. As an indirect proxy for neural activity, the BOLD signal primarily reflects hemodynamic responses including changes in blood flow and oxygenation, and therefore reflects only a single dimension of brain function. Accumulating evidence indicates that hemodynamic signals can dissociate from metabolic activity under various physiological and experimental conditions [26–29], indicating that there are metabolic dimensions of brain function that are not directly accessible with BOLD-fMRI. Moreover, BOLD signals are susceptible to systemic physiological confounds — such as fluctuations in end-tidal CO2 [30], the volume of red blood cells in the blood [31], respiratory volume [32], and heart rate [33]. Therefore, while powerful, BOLD-fMRI provides only a partial view of cortical functional architecture when considered in isolation, particularly with respect to underlying metabolic processes.

Recent advances in positron emission tomography (PET) technology, including increased scanner sensitivity and the adoption of constant-infusion protocols, have enabled the tracking of intra-scan dynamic changes in cerebral glucose metabolism with [^18^F]-fluorodeoxyglucose (FDG). Unlike traditional bolus injections, constant-infusion maintains a more stable plasma concentration of FDG and enables more sensitive measurement of fluctuations in glucose utilization [34,35]. Building on these methodological developments, functional FDG-PET (fPET-FDG) has emerged as a promising technique for capturing time-resolved metabolic responses in the human brain, thereby attracting growing interest in the field [36,37]. A growing body of work now demonstrates that fPET-FDG can effectively map dynamic metabolic responses elicited by sensory and higher-order cognitive processes [27,38–41].

In light of the multidimensional physiological basis of brain organization, there is increasing interest in using fPET-FDG to characterize synchronous glucodynamic fluctuations across the brain regions, referred to as resting-state metabolic connectivity (RSMC) in the literature [28,42–44]. Previous RSMC studies revealed a dimension of brain organization that complements, yet diverges from, functional architecture inferred from hemodynamics alone. In particular, metabolic connectivity patterns show only moderate correspondence with canonical fMRI networks and uncover unique features, such as prominent frontoparietal connectivity patterns, not readily observed when using fMRI [28,45–48]. Furthermore, RSMC appears to capture distinct aspects of cognitive function and aging that extend beyond the insights provided by fMRI-based functional connectivity [49–51]. Collectively, these findings indicate that by capturing and tracking regional dynamics of glucose consumption — the primary fuel of brain function — metabolic connectivity reflects meaningful organizational properties of the brain that extend beyond, and complement, hemodynamic measures.

Despite these advances, existing RSMC studies have largely emphasized exploratory analyses of large-scale networks, leaving the spatiotemporal properties of metabolic connectivity insufficiently characterized. How metabolic connectivity is organized across the cortex at both global and local scales, and how this organization relates to established anatomical, functional, and metabolic principles, remains largely under explored. In addition, although advances in PET instrumentation and image reconstruction now permit fPET data reconstruction at temporal resolutions of only a few seconds [52–54], FDG phosphorylation is inherently slow. As a result, it remains unclear what intrinsic temporal properties, or effective “speed”, of glucodynamic interactions are captured by fPET-FDG-derived RSMC.

In this study, we aimed to provide a systematic evaluation of RSMC by characterizing its 1) topological profiles across spatial scales, 2) temporal properties, and 3) relationships to other features of brain organization. Using a publicly available simultaneous fPET-fMRI dataset (Monash rsPET–MR) [52], we delineated metabolic organization across spatial scales by connectivity-based boundary mapping [17–20], network community detection [55,56], and principal gradient analysis [57,58]. We further filtered the fPET-FDG signal into distinct temporal frequency bands to identify the temporal components that dominate metabolic connectivity. Finally, we contextualized these RSMC spatial patterns within known anatomical, functional, and energetic constraints.

Note that for clarity and consistency with emerging nomenclature conventions [43,44], we refer to temporal correlations derived from BOLD-fMRI time series as “functional connectivity” and those derived from fPET-FDG time-activity curves (TACs) as “metabolic connectivity”. We acknowledge that BOLD signals are closely linked to oxygen “metabolism”, and glucose metabolism — as measured by fPET-FDG — is tightly coupled to dynamic changes in brain “function”. A preliminary account of this study has been presented in abstract form [59].

## Materials and Methods

### Dataset and acquisition protocol

We utilized a publicly available resting-state functional PET–MRI dataset [52], which consists of simultaneous fPET-FDG and BOLD-fMRI scans from 27 healthy young adults (aged 18–23 years, mean age 19 years). Data were acquired on a Siemens 3T Biograph mMR scanner during a 95-minute scan session, and the subjects were instructed to rest with their eyes open throughout the experiment. FDG was continuously infused at an average dose of 233 MBq and at a rate of 36 mL/h. One participant was excluded from our study due to an infusion pump error, and two participants were excluded due to abnormal whole brain tracer kinetics, resulting in a sample size of 24 subjects, as in our previous study [60].

Functional PET-FDG data were acquired in list mode, and initiated simultaneously with the onset of FDG radiotracer infusion. Attenuation correction for fPET-FDG data was performed using an Ultrashort TE (UTE) MRI, and the PET data were reconstructed using the Ordinary Poisson Ordered Subset Expectation Maximization (OP-OSEM) algorithm with a nominal voxel size of 2.09 × 2.09 × 2.03 mm^3^ and a temporal resolution of 16 seconds per frame [52].

During the first 30 minutes of the scan session, data from several non-fMRI pulse sequences were acquired while allowing the radiation of PET tracer to gradually increase to detectable levels, including a high-resolution T1-weighted MPRAGE (total acquisition time = 7.01 min, TR = 1640 ms, TE = 2.34 ms, flip angle = 8°, field of view (FOV) = 256 × 256 mm^2^, voxel size = 1 × 1 × 1 mm^3^, 176 sagittal slices). At the 30-minute time point, six consecutive 10-minute BOLD-fMRI scans were performed using T2*-weighted echo-planar imaging (TR = 2450 ms, TE = 30 ms, FOV = 190 mm, voxel size = 3 × 3 × 3 mm^3^, 44 axial slices acquired in ascending interleaved order).

### Preprocessing

An overview of the data preprocessing pipeline is shown in **Figure 1**.

**Figure 1.**
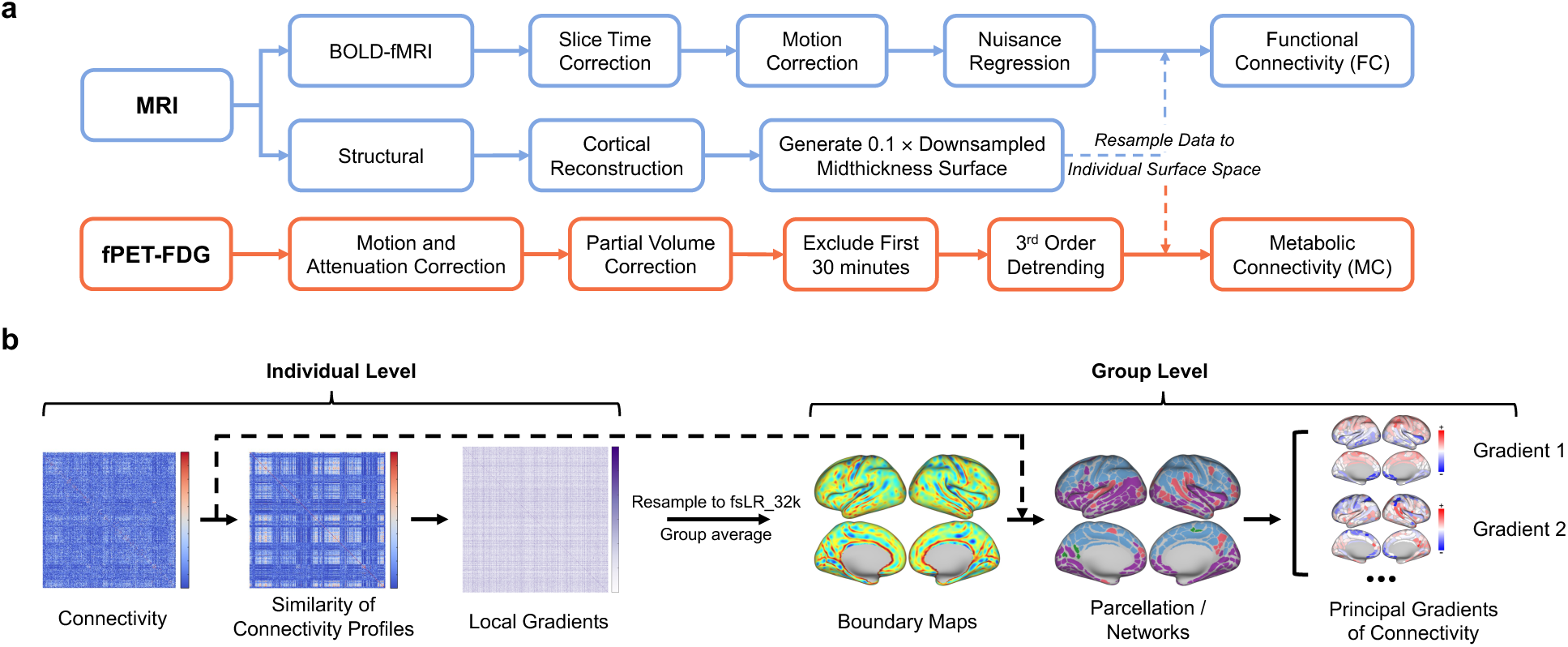
Overview of data preprocessing and analysis pipelines: (a) Preprocessing: BOLD-fMRI data underwent slice timing correction, motion correction, and aCompCor-based nuisance regression. fPET-FDG data were corrected for motion, attenuation, and partial volume effects; the first 30 minutes of data were discarded to match the total scan duration of resting-state BOLD-fMRI acquisitions, followed by a third-order polynomial detrending to remove the low-frequency baseline of FDG accumulation. Structural MRI data was processed using FreeSurfer’s recon-all, and a midthickness surface downsampled to 1/10 of original vertices was generated for each subject. Both BOLD-fMRI and fPET-FDG data were resampled to this individual midthickness surface for ensuing analysis. (b) Connectivity-based analysis: Resting-state functional and metabolic connectivity matrices were transformed into vertex-wise connectivity correlation maps, from which local gradient maps were computed for each subject. These gradients were used to generate group-level boundary maps and delineate fine-grained cortical parcels. The resulting cortical parcellation was further used to identify parcel-wise network communities and to extract the principal gradients of connectivity.

#### Functional PET-FDG preprocessing

Preprocessing of fPET-FDG data included the following steps: (1) motion correction using AFNI [61], (2) gray-matter partial-volume correction using PETSurfer [62], (3) exclusion of the first 30 min after tracer infusion onset, to better approximate a steady state of FDG kinetics (i.e., a relatively constant plasma FDG concentration) and to align the fPET-FDG data with the BOLD-fMRI acquisition window, and (4) temporal detrending to mitigate the effects of slow baseline FDG accumulation over time. To evaluate the impact of different preprocessing strategies, we tested three detrending methods (second-order polynomial, third-order polynomial, and global signal regression (GSR)), with or without the initial 30-min period. Unless otherwise specified, all main analyses were based on third-order detrending without the initial 30-min period.

#### BOLD-fMRI preprocessing

Preprocessing of BOLD-fMRI data included the following steps: (1) slice timing correction using AFNI [61], (2) rigid-body motion correction using AFNI, and (3) nuisance regression with the aCompCor method to remove confounding physiological fluctuations and enhance the neuronal specificity of functional connectivity measurements [63]. White matter and ventricular masks were derived from high-resolution structural MRI data using FreeSurfer [64] and the first five principal components of all white-matter and ventricular signals were included as nuisance regressors. We also tested preprocessing using GSR or without nuisance regression to see their influence on RSFC boundary mapping.

#### T1-weighted structural data preprocessing

We automatically reconstructed cortical surfaces from the T1-weighted structural data using FreeSurfer’s *recon-all* pipeline [64], and from them generated the mid-thickness surfaces and downsampled the resulting surfaces to 10% of the original numbers of vertices to reduce computational complexity and mitigate artifacts that may arise from volume-to-surface resampling and uneven vertex spacing of meshes [65,66]. The preprocessed BOLD-fMRI and fPET-FDG data were coregistered and resampled onto the downsampled mid-thickness surface for each subject. We then applied surface-based smoothing using a 10-mm full-width at half-maximum (FWHM) Gaussian kernel to the fPET-FDG data to reduce noise and improve sensitivity, while the BOLD-fMRI data were left unsmoothed given their higher signal-to-noise ratio (SNR).

### Connectivity-based boundary mapping

We followed the connectivity-based boundary mapping framework proposed by Gordon *et al.* [20]. For each subject, we estimated RSFC and RSMC as Fisher Z-transformed Pearson correlations between all vertex pairs, then constructed connectivity similarity matrices by correlating the connectivity profiles across all pairs of vertices. We then derived local gradients from these connectivity similarity matrices on each subject’s mid-thickness surface using the *cifti-gradient* function in Connectome Workbench [67], thereby capturing abrupt spatial transitions in connectivity similarity.

To reduce spurious correlations introduced by volume-to-surface mapping and uneven vertex spacing of meshes, which can be especially prominent under low signal-to-noise conditions [65,66], we used a local gradient regression approach (see **Figure 2a**). For each subject, we generated surrogate datasets by randomly shuffling the time series at each surface vertex. This procedure preserves the temporal structure of each vertex’s signal (e.g., its power spectrum) while removing temporal correlations between vertices. The surrogate datasets were then processed using the same preprocessing and analysis pipeline as the empirical data. Because the surrogate data contain no true correlations between vertices, local gradients estimated from these data reflect effects driven by volume-to-surface resampling and smoothing on unevenly sampled meshes rather than genuine metabolic connectivity. These surrogate-derived gradients were then linearly regressed out of the corresponding empirical gradients, thereby reducing the influence of these confounding factors. To examine how confounding factors and correction via local gradient regression affect the boundary mapping results, we created synthetic datasets at different SNR levels for validation. Further methodological details are provided in Error! Reference source not found..

**Figure 2.**
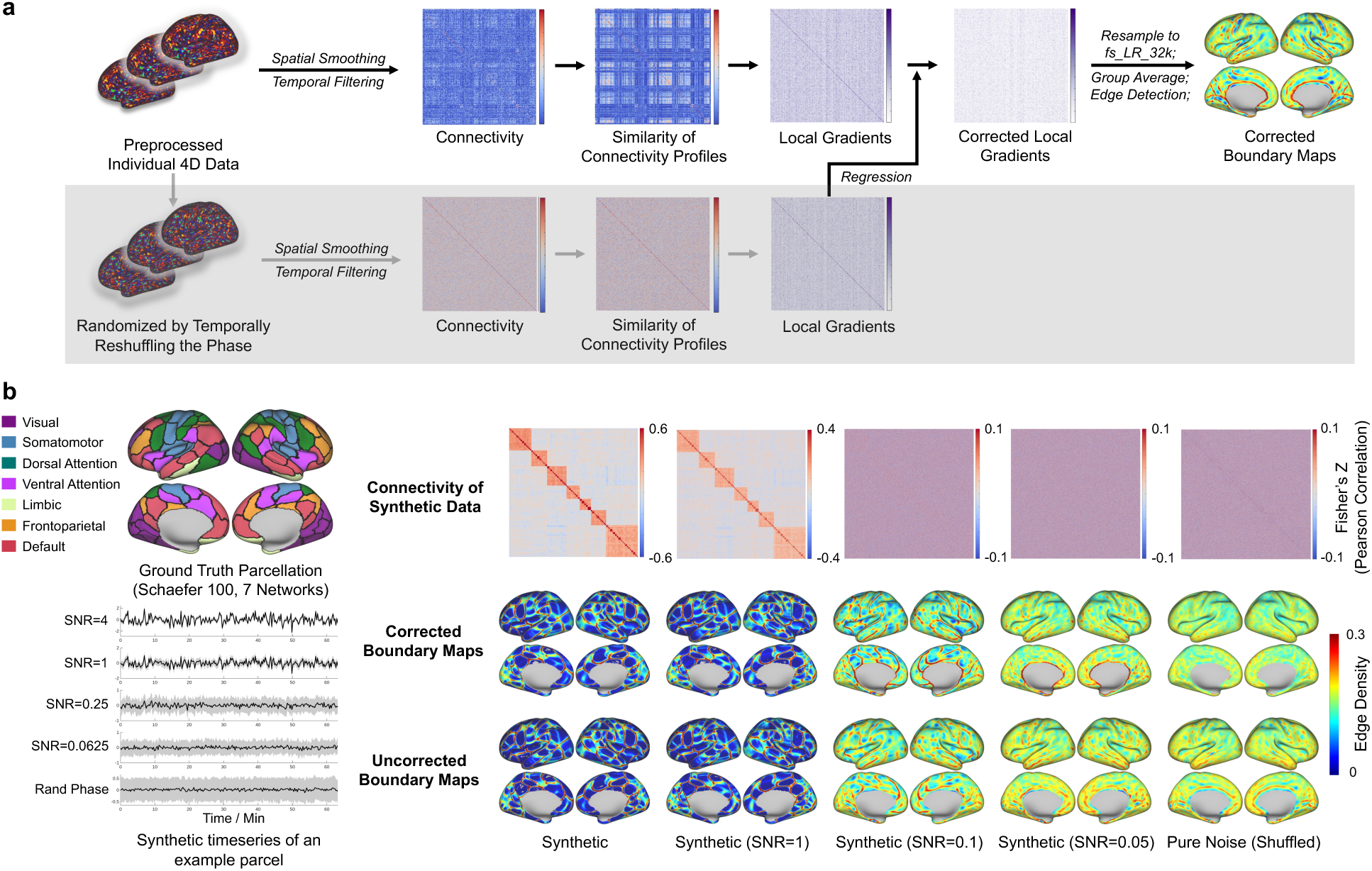
Overview of the connectivity-based boundary mapping strategy and its validation using synthetic data. (a) The framework for characterizing and correcting structured noise arising from volume-to-surface resampling and meshes with unevenly spaced vertices, using a control dataset generated by randomly reshuffling the phases of each individual’s empirical time-series data. (b) Evaluation of the proposed boundary mapping approach using synthetic datasetswith varying SNR levels. Left: Synthetic “ground-truth” data based on the Schaefer 7-network, 100-parcel atlas, with exemplar time series shown for a representative parcel, with voxel-wise variability depicted as shading. Right: Connectivity matrices and boundary maps, with and without correction for structured noise.

The corrected local gradients were then resampled to the common surface space (fsLR 32k [68]), averaged across subjects, and smoothed with a 6-mm FWHM Gaussian kernel. A watershed-by-flooding algorithm [20,69] was then applied to the local gradients of each vertex, delineating regions of abrupt transitions in connectivity similarity across the cortical surface. Boundary probability maps (hereafter referred to as boundary maps) were then obtained by averaging the boundaries derived from the gradients of each vertex, representing the likelihood that each cortical vertex lies on an areal boundary. Cortical parcels were additionally derived from the group-level boundary map following the approach described by Gordon *et al.* [20] to define regions for network detection.

### Network detection

To further explore the global organization of the RSFC and RSMC, we used the parcels derived from boundary mapping to compute parcel-wise connectivity. We applied the Louvain community detection method [55] from the Brain Connectivity Toolbox [56] to parcel-wise connectivity to identify cortical networks. This method partitions the brain into distinct functional networks (or communities) by optimizing modularity, ensuring that regions within the same network exhibit stronger connectivity compared to those from different networks. To obtain network partitions with a target number of communities (*k* = 2–8), we iteratively adjusted the resolution parameter 𝛾, with higher 𝛾 encouraging more networks to be detected. Starting from an initial 𝛾 = 1, the community detection algorithm was repeatedly applied. If the resulting partition contained fewer communities than the target *k*, 𝛾 was increased by 0.05 to promote finer-grained partitions, and vice versa. Only runs in which the detected number of communities exactly matched the target *k* were retained and counted as valid iterations. We set the maximum number of iterations to 100,000 and allowed up to 200 rounds for the algorithm to converge to the target number of networks.

To test whether the identified network structure exceeded chance, we computed the mean silhouette value of the parcel-wise connectivity for both RSFC and RSMC. For each network, a null distribution of the observed silhouette value was constructed by computing silhouette values on sham parcellations, generated by randomly permuting the labels of the community assignment (2000 iterations) while fixing the overall connectivity matrix. Two-sided nonparametric p-values of the empirical silhouette values were subsequently obtained according to the null distribution.

### Principal gradients of connectivity

In order to characterize the continuous global organization of cortical metabolic connectivity, we examined the principal gradients of RSFC and RSMC (hereafter referred as “functional gradients” and “metabolic gradients”). Diffusion map embedding [57] implemented in the BrainSpace toolbox [58] was applied to the parcellated, group-averaged connectivity matrices. In this framework, the diffusion embedding identifies low-dimensional manifolds that best preserves the similarity structure of the input connectivity matrix, with the leading eigenvectors (gradients) capturing the dominant axes of connectivity variation. The resulting functional and metabolic gradients thus represent continuous axes of connectivity variation, capturing gradual, large-scale transitions across the cortex. All analyses used BrainSpace default parameters, with the input connectivity matrices used as affinity matrices. To assess the robustness of gradient estimates to parcellation choice and to facilitate comparison with other organizational metrics, principal gradients were computed using three different cortical parcellations: a study-specific custom parcellation (386 regions of interest (ROIs) for RSMC and 403 ROIs for RSFC), the Glasser atlas (360 ROIs) [70], and the Schaefer atlas (400 ROIs) [71].

### Frequency-dependent analysis of RSMC

To examine the temporal frequency dependence of RSMC and its spatial organization, several frequency-filtered variants of the fPET-FDG data were generated. Sixth-order Butterworth low-pass (< 1/48 Hz, 1/64 Hz, 1/128 Hz, 1/256 Hz) or high-pass (> 1/48 Hz, 1/ 64 Hz, 1/128 Hz, 1/256 Hz) temporal filtering was applied to each vertex to isolate different frequency components.

Each filtered dataset was then processed with the same analytical pipeline as in the main analysis, including RSMC estimation, boundary mapping, and principal gradient extraction. This design allowed us to assess how different temporal frequency components influence RSMC and its associated spatial patterns.

### Correlating metabolic gradients to anatomical, functional, and energetic measures

To facilitate interpretation of the large-scale axes captured by metabolic gradients, we examined their spatial correspondence with a set of previously established anatomical, functional, and energetic metrics.

Anatomical measures included principal gradients of microstructural covariance derived from cortical thickness [72], eigenmodes of cortical surface geometry [73], smoothed cortical myelin map [68], and geodesic distance to paleocortex [74].

Functional measures comprised RSFC functional gradients estimated within the present dataset [52], as well as multifaceted functional gradients that integrate multiple spontaneous fluctuation features (such as power of low-frequency fluctuations and regional homogeneity) and network properties (degree centrality, path length, global and local efficiency) derived from BOLD-fMRI [74].

Energetic measures consisted of cerebral metabolic rate of glucose (CMRGlu) derived from the present dataset [52], together with mean cortical gene expression profiles linked to core energy metabolism pathways, including the pentose phosphate pathway (ppp), glycolysis, the tricarboxylic acid cycle (tca), oxidative phosphorylation (oxphos), and lactate metabolism [75].

All reference metrics were parcellated using either the Schaefer 400-ROI atlas [71] or the Glasser 360-ROI atlas [70], consistent with the parcellation used for metabolic gradient analyses. Spatial correspondence between metabolic gradients and each reference measure was quantified using Pearson correlation coefficients across cortical parcels. Statistical significance was assessed using corresponding two-tailed p-values.

## Results

### Impact of different preprocessing on connectivity

For RSMC, different detrending strategies had a substantial impact on the spatial structure of metabolic connectivity, as also discussed by Coursey *et al.* [60]. As shown in **Figure S1**, second-order polynomial detrending left visible structured background correlations, which were effectively suppressed by third-order detrending. Although GSR further reduced background correlations, it also introduced artificial anticorrelations. Excluding the initial 30 minutes of data attenuated structured background correlations across all three detrending approaches, consistent with the expectation that tracer kinetics had not yet approximated pseudo steady-state during this early phase. Accordingly, third-order detrending combined with exclusion of the first 30 minutes was adopted for all subsequent RSMC analyses.

For RSFC, boundary maps derived from different nuisance regression strategies were highly consistent (**Figure S2**). Spatial boundary organization was well preserved regardless of whether aCompCor [63] or GSR was applied, with only minor differences compared to maps generated without nuisance regression. Given the ongoing debate regarding GSR and its potential to introduce artificial anticorrelations [76,77], subsequent RSFC analyses were conducted using data processed with the aCompCor approach.

### Local metabolic organization revealed by boundary mapping

Mapping local boundaries from RSMC is particularly challenging due to the intrinsically lower SNR of fPET-FDG data, and is made possible by a local gradient regression approach in our study. As shown in **Figure S3**, volume-to-surface resampling and boundary mapping performed without correction introduced structured boundary-like patterns even in the absence of true connectivity structure, potentially driven by cortical curvature. Analyses on synthetic datasets demonstrated that under low-SNR conditions, these confounding factors can dominate the resulting boundary maps, obscuring genuine local transitions (**Figure 2b**). Gradient regression effectively reduced artifact contamination under low SNR, enabling partial recovery of meaningful connectivity boundaries while leaving high-SNR conditions largely unaffected (**Figure 2b**, see also **Figure S4 and Supplementary Methods**). The combined application of local gradient regression and surface-based smoothing of timeseries further stabilized boundary estimates and reduced curvature-like artifacts (**Figure S5**).

As illustrated in **Figure 3a**, boundary maps derived from RSMC after gradient regression correction revealed discernible local transitions across the cortical surface. Although these metabolic boundaries did not systematically align with areal borders derived from RSFC, prominent edges were observed in several regions, including the primary and secondary visual cortices and the primary somatosensory cortex. These transitions suggest that adjacent cortical areas in these regions exhibit distinct metabolic connectivity profiles, indicative of localized metabolic organization.

**Figure 3.**
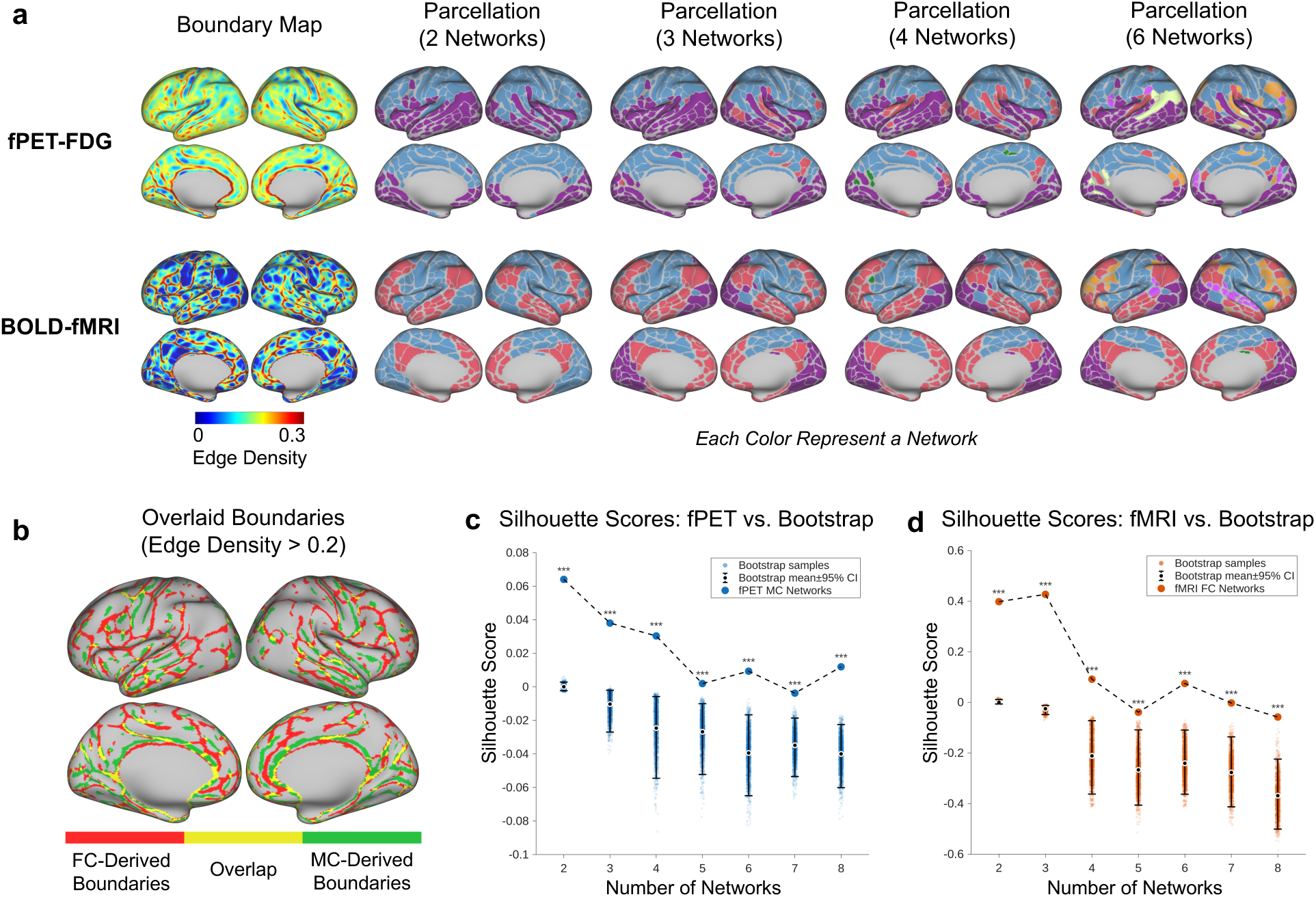
Boundary maps and cortical network organization: RSMC vs. RSFC. (a) Group-level boundary maps, cortical parcellations, and networks identified based on MC and FC respectively. (b) Overlaid RSFC and RSMC boundary maps. Vertices with edge density (probability) larger than 0.2 are presented. Within-network homogeneity of (c) RSMC- and (d) RSFC-derived parcellations represented by silhouette scores averaged across all cortical parcels, tested against bootstrap samples with randomly shuffled network assignment.

In contrast, boundary maps derived from RSFC were more spatially distinct and structured, with a higher density of boundaries that clearly delineated well-established cortical divisions, such as primary somatosensory cortices, precuneus and posterior cingulate cortex, consistent with prior RSFC-based boundary mapping studies [18–20]. Direct comparison revealed partial spatial overlap between RSFC- and RSMC-derived boundaries (**Figure 3b**), indicating that certain local organizational features could be shared across metabolic and hemodynamic connectivity.

### Global metabolic organization from network detection and principal gradients

#### Superior-inferior differentiation in RSMC networks

To assess the global organization of RSMC, we estimated cortical networks using the Louvain community detection algorithm based on the parcels derived from boundary maps (**Figure 3a**). Across different network resolutions, RSMC exhibited a clear differentiation between frontal-parietal (superior) and temporal-occipital (inferior) regions, whereas RSFC reproduced canonical resting-state networks, including the default mode, visual, and frontoparietal systems [20–24]. Although RSMC networks exhibited overall lower silhouette scores than RSFC, the values remained significantly above chance (*p* < 0.001), indicating the presence of meaningful network organization (**Figure 3c, d**, also see **Figure S6** for RSMC matrices reordered according to superior-inferior network delineation).

#### Homotopic connectivities in RSMC networks

Similar to RSFC, homotopic connectivity patterns were visible in the RSMC matrices (**Figure 4a**), with similar connectivity profiles between corresponding regions of the left and right hemispheres. This pattern was also apparent in the low-pass filtered data (**Figure S7**).

**Figure 4.**
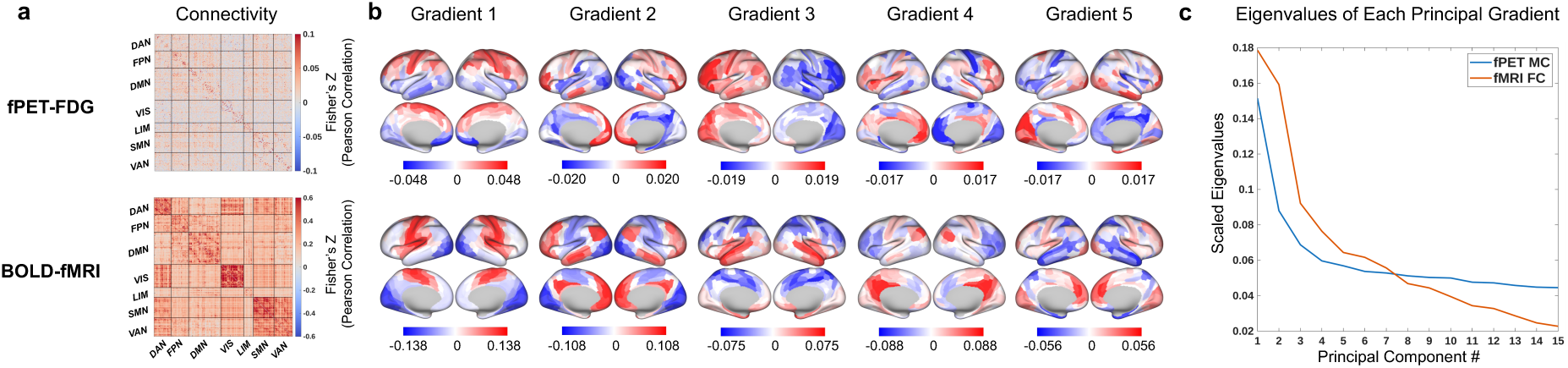
Connectivity and principal gradients: RSMC vs. RSFC. (a) Group-level RSMC and RSFC matrices parcellated in Glasser 360-ROI atlas, and (b) principal gradients derived from RSMC and RSFC matrices. (c) First 15 eigenvalues (scaled to a sum of 1) representing variance explained by each functional gradient and metabolic gradient.

#### Principal gradients of metabolic network organization

Principal gradients exhibited continuous axes of cortical organization (**Figure 4b**). The first metabolic gradient (hereafter referred as metabolic G1) revealed a prominent superior-inferior axis, aligning with the major network-level division observed in RSMC, whereas the second metabolic gradient (hereafter referred as metabolic G2) captured a weaker anterior-posterior differentiation. In contrast, RSFC functional gradients reproduced the well-established unimodal-to-transmodal hierarchy described by Margulies *et al.* [25]. The steeper eigenvalue decay observed for metabolic gradients (**Figure 4c**) indicates that the first two components represent the dominant modes of metabolic connectivity.

Notably, across different detrending strategies and data exclusion schemes in fPET, we consistently observed a superior-inferior metabolic gradient in RSMC (**Figure S8**). Comparable large-scale organization was also evident across different cortical parcellations (**Figure S9**), confirming the robustness of the characterized superior-inferior axis.

### Low-frequency components predominate in metabolic networks and gradients

To investigate how the spectral properties of fPET dynamics shape overall connectivity organization, the signals were separated into low- and high-frequency (cutoff at 1/64 Hz) components. Low-frequency components largely preserved the connectivity profiles and the first two metabolic gradients observed in the unfiltered data (**Figure 5a, b**), whereas high-frequency components yielded markedly weaker spatial organization and resembled the randomly-shuffled control, indicating a reduced contribution to the systemic RSMC structure.

**Figure 5.**
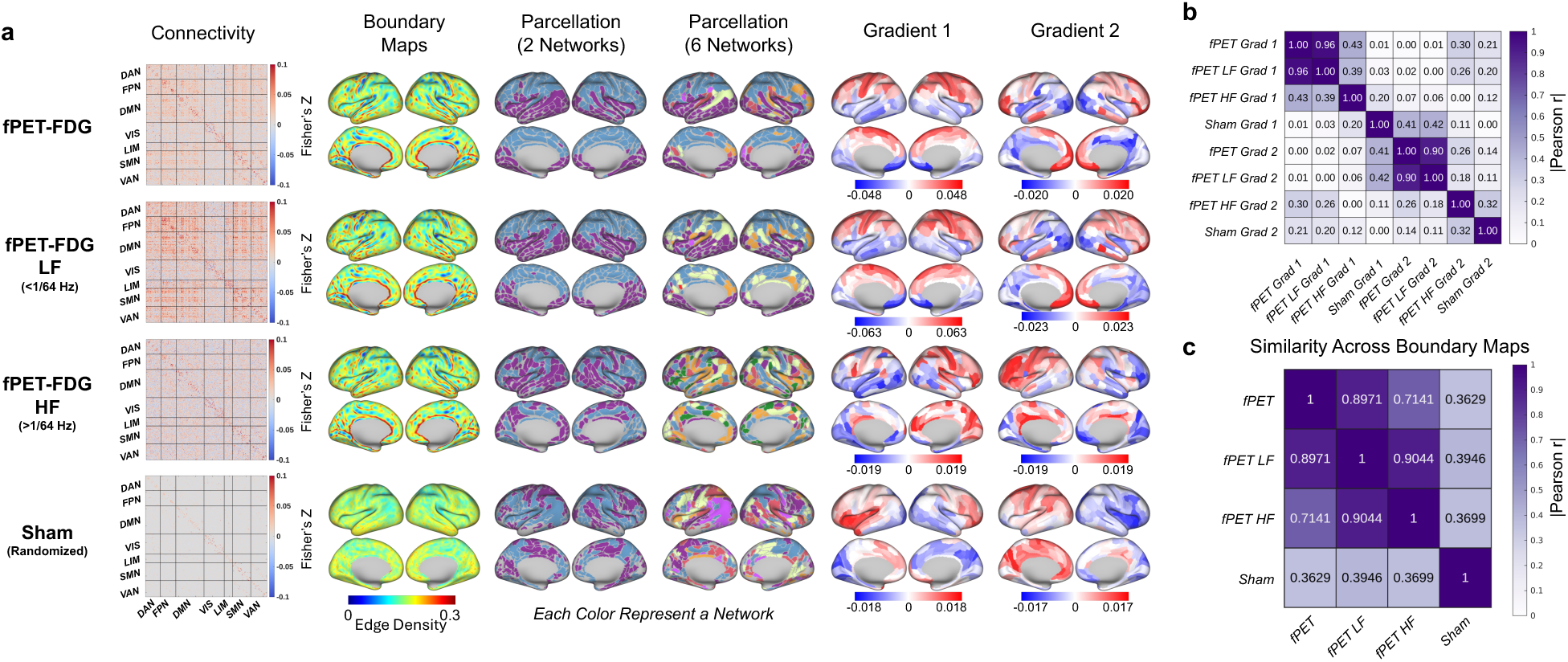
Low-frequency components dominate RSMC network features, while both low-frequency (LF) and high-frequency (HF) components shape the boundary map. (a) Connectivity matrices, boundaries, parcellations, and principal gradients derived from RSMC, both with and without temporal filtering. A sham dataset with phase-randomized timeseries that went through the same processing steps is shown for reference. (b) Pearson correlation between principal gradients estimated across fPET-FDG datasets. The first two principal gradients derived from low frequency component closely resembled those derived from unfiltered data, and diverge from those derived from high frequency component or sham data. (c) Pearson correlation between boundary maps estimated across fPET-FDG datasets. Boundary maps from both low- and high-frequency bands resembled the unfiltered data and diverged from the sham.

Nevertheless, both frequency ranges contributed to local boundary formation: low-frequency dynamics dominated global network architecture and gradient structure, whereas high-frequency components retained localized boundary features beyond those observed in the sham control (**Figure 5c**). These results suggest that slow fluctuations in fPET-FDG dynamics primarily drive the observed large-scale metabolic organization, while both slow and faster dynamics influence local boundary transitions.

To further examine the contribution of different frequency components, we evaluated metabolic connectivity across a broader set of predefined frequency bands. As shown in **Figure S7**, the dominant patterns of metabolic connectivity (strong frontoparietal connectivity) remained visible even under extensive low-pass filtering (< 1/256 𝐻𝑧, retaining approximately one-eighth of the original frequency spectrum). This persistence suggests that slow FDG dynamics make a substantial contribution to the global RSMC and its spatial organization.

### Relating metabolic gradients to anatomical, functional, and energetic constraints

Correlation analyses revealed several anatomical, functional, and energetic measures that showed moderate-to-strong spatial correspondence with the metabolic gradients (**Figure 6**). For metabolic G1, which captured the most prominent superior-inferior axis, we observed significant correlations with the second principal gradient of microstructural covariance derived from cortical thickness [72] (𝑟 = 0.5873, 𝑝 < 10^−3^), the third eigenmode of cortical surface geometry [73] (𝑟 = 0.6355, 𝑝 < 10^−3^), and geodesic distance to paleocortex [74] (𝑟 = 0.5262, 𝑝 < 10^−3^). In addition, G1 showed a significant negative correlation with the multifaceted functional gradient derived from BOLD-fMRI [74] (𝑟 = −0.4812, 𝑝 < 10^−3^). Among energetic measures, G1 was significantly correlated with the mean cortical gene expression associated with the tca pathway (𝑟 = 0.5306, 𝑝 < 10^−3^) and the lactate metabolic pathway [75] (𝑟 = 0.5200, 𝑝 < 10^−3^).

**Figure 6.**
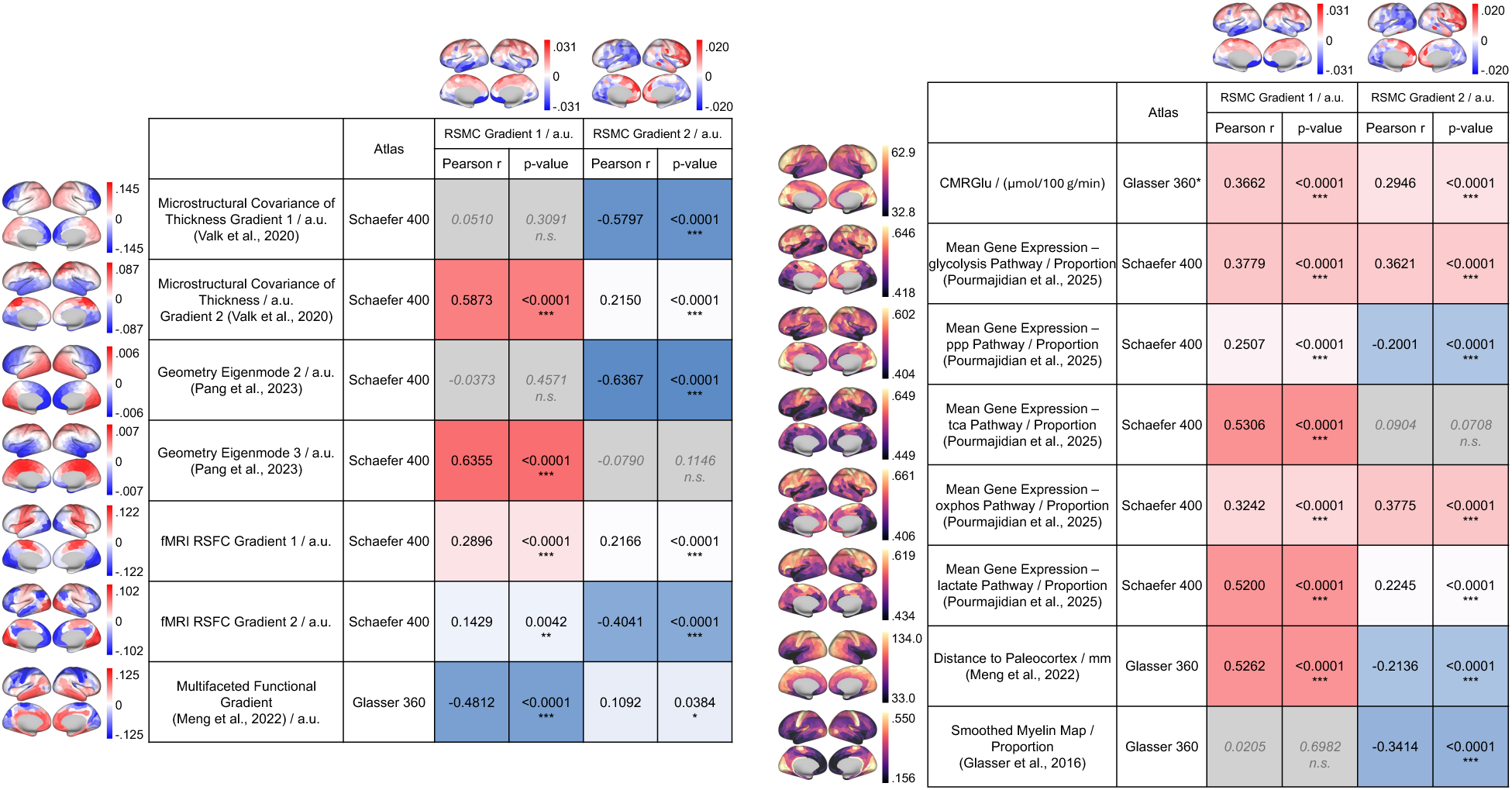
Spatial similarity between metabolic gradients and diverse macroscale cortical features spanning structure, function, metabolism, and evolution. Gradient features (left table) are shown using a red-white-blue colormap, and other non-negative features (right table) are shown using a magma colormap. Red and blue colors in the table indicate positive and negative Pearson correlations between metabolic gradients and reference measures, respectively, with intensity indicating the absolute correlation strength. Metrics not significantly correlated with metabolic gradients are colored in gray. Metrics significantly correlated to RSMC gradients are labeled with asterisks (*, **, ***: 𝑝 < 0.05, 0.01, 0.001, respectively). The RSMC metabolic gradients shown here above the table were derived in current study, rendered on Schaefer 400-ROI atlas for comparison. For metrics using Glasser 360-ROI atlas, comparison are conducted with RSMC metabolic gradients parcellated in Glasser 360-ROI atlas. Refer to Figure S9 for metabolic gradients rendered on different parcellation schemes.

For the anterior-posterior axis revealed by metabolic G2, we observed significant negative associations with the first principal gradient of microstructural covariance [72] (𝑟 = −0.5797, 𝑝 < 10^−3^) and the second eigenmode of cortical surface geometry [73] (𝑟 = −0.6367, 𝑝 < 10^−3^). In the functional domain, G2 also showed a moderate negative correlation with the second RSFC functional gradient derived from BOLD-fMRI (𝑟 = −0.4041, 𝑝 = 10^−3^).

## Discussion

In this study, we systematically examined the spatiotemporal organization of cortical metabolic connectivity using fPET-FDG-derived RSMC. By characterizing metabolic organization across spatial scales, we identified both global architectural features and local metabolic transitions that are not readily captured by hemodynamic measures alone. Extending prior studies that evaluated metabolic artchitecture through simplified nodal connectivity matrices, our analyses revealed a dominant superior-inferior organization separating frontoparietal and temporo-occipital regions, along with a secondary anterior-posterior axis reflected in the second principal metabolic gradient. The dominant axis was consistently observed across network-based analyses and was robust to different detrending approaches. Importantly, frequency-resolved analyses indicate that global, long-range RSMC architecture is predominantly shaped by slow, minute-scale FDG-based glucodynamic fluctuations. At the same time, both slower and relatively faster temporal components contribute to the local metabolic organization, as reflected in RSMC-derived boundary maps. Together, these findings offer new insights into brain functional organization from a metabolic perspective.

### Spatial organization: RSFC vs. RSMC

In agreement with previous research focusing on simplified matrix representations of brain connectivity [28,50,51], the metabolic network topography characterized here via fPET-FDG notably departs from canonical RSFC networks. This divergence corroborates the existence of distinct organizational principles underlying glucodynamics and hemodynamics. Such discrepancies between RSMC and RSFC likely arise from a combination of physiological and technical factors, as noted in previous studies [28]. Biologically, RSMC and RSFC represent disparate facets of brain activity: fPET-FDG tracks quasi-steady-state glucose utilization, whereas BOLD-fMRI captures transient vascular and blood-oxygenation fluctuations. This physiological decoupling is evidenced by known dissociations between blood flow and glucose metabolism during conditions such as neural inhibition [26,78] and systemic physiological challenges [35], as well as oxygen and glucose metabolism uncoupling in the presence of aerobic glycolysis [79,80]. Furthermore, glucose metabolism underlying fPET operates on a significantly slower temporal scale, constrained by the steady-state requirements of FDG kinetics, in contrast with second-scale hemodynamic signals. Consequently, RSMC and RSFC may capture the brain connectome across distinct timescales. It is also noteworthy that the inherently lower sensitivity of fPET-FDG may limit the detection of subtle metabolic connectivity patterns at current stage. As such, part of the observed divergence between RSFC and RSMC may reflect technical constraints rather than purely physiological differences, and future studies with larger dataset and improved sensitivity may reveal additional details.

### Temporal characteristics of RSMC

With ongoing advances in PET technology, it has become feasible to reconstruct short-frame fPET-FDG data with sufficient quality, evolving from early protocols using minute-long frames to more recent methods achieving frame durations down to second-scale sampling intervals [52–54]. However, our results suggest that long-range metabolic connectivity — that drives metabolic network delineations — is primarily shaped by minute-scale, low-frequency fluctuations of the fPET-FDG signal. These observations suggest that rapid, second-scale image reconstructions may not be needed for resolving coherent fPET fluctuations giving rise to long-range metabolic connectivity. One possible explanation lies in the intrinsic low-pass filtering property of FDG tracer kinetics — governed by the rate of tracer exchange between plasma and tissue, and by tracer accumulation — which inherently attenuates fast fluctuations while amplifying slower trends. From a technical perspective, current fPET-FDG acquisition and reconstruction methods may also yield limited SNR at sub-minute frequencies, limiting our ability to resolve the metabolic organization encoded in faster signal components. Another plausible factor is the contribution of minute-scale variations in neuronal states, such as arousal fluctuations or infra-slow brain activity that modulates cortical excitability and cerebral metabolism [81]. Given the 60-minute scan duration, these global state changes likely exert a substantial influence on the observed metabolic patterns. This hypothesis could be further clarified by correlating fPET-FDG fluctuations with concurrent EEG or pupil-diameter measurements in future studies.

### Anatomical, functional and energetic correlates of the RSMC principal gradients

As shown in **Figure 6**, we examined how RSMC organization relates to anatomical, functional, and metabolic metrics to explore the physiological context of metabolic organization.

The metabolic G1 identified in this study exhibits a prominent superior-inferior axis, a pattern that was consistently recovered across both gradient- and network-based analyses. This axis is not unique to RSMC, convergent superior-inferior organizational patterns have been reported across multiple domains, including geometrical eigenmodes of the cortical surface [67], principal gradients of cortical thickness covariance [66], multifaceted functional gradients [74], and spatial maps of gene expression related to energy metabolism [75], particularly genes involved in the TCA cycle and lactate pathways, which are thought to reflect core, developmentally established metabolic programs in the healthy brain. A parsimonious interpretation is that these convergent axes reflect shared metabolic demand constraints underlying cortical geometry, structural differentiation, and macroscale functional properties.

On the other hand, the superior-inferior axis of metabolic G1 has also shown a clear correspondence with distance to the paleocortex (adapted from Meng *et al.* [74]), aligning with the canonical dorsal-ventral dichotomies as described by the brain’s dual-origin theory[82–85]. Within this theoretical framework, cortical organization is shaped by two broad developmental trajectories emerging from paleocortical and archicortical origins, giving rise to a dorsal-ventral organizational dichotomy of cortical systems. These systems are thought to support different classes of functional specialization, with dorsal regions preferentially associated with spatial, temporal, and action-related processes, and ventral regions more closely linked to semantic, affective, and motivational functions [82–85]. The observed alignment between metabolic G1 and paleocortical distance, and also other anatomical, functional and metabolic measures raises the possibility that shared developmental origins contribute to long-range similarities across various domains, leading to the consistent emergence of a superior-inferior axis across multiple modalities. Notably, previous work on cortical thickness covariance [72] and multifaceted functional gradients has similarly proposed that large-scale superior-inferior axes may reflect the dual developmental origins of the cerebral cortex.

A second, anterior-posterior axis represented by G2 showed a broadly similar relationship with the first cortical thickness gradient and the second geometrical eigenmode. Although the anterior-posterior axis was less consistent across parcellation schemes (**Figure S9**), it may still reflect meaningful variation in how metabolic organization relates to underlying microstructural and geometrical features. Further works with larger dataset with increased SNR are needed to evaluate the robustness and interpretation of this secondary gradient.

### Limitations and future directions

The findings of this study are subject to a few considerations that warrant further investigation in future work. First, although the sham data-based gradient regression approach helps mitigate artifactual connectivity gradients arising from anatomical geometry and registration, it relies on the implicit assumption that artifacts introduced by volume-to-surface resampling and smoothing on unevenly sampled meshes are statistically orthogonal to genuine functional or metabolic gradients. Under this assumption, regressing out sham-derived patterns would not distort intrinsic functional or metabolic connectivity patterns. The largely preserved network delineations observed in fMRI-based functional connectivity following regression provide some support for this premise; however, the impact of this regression on the spatial organization of fPET-based metabolic connectivity remains to be systematically evaluated. Importantly, this potential limitation is more likely to affect the precision of specific areal boundaries than the broader network organization. As PET instrumentation continues to evolve, offering sensitivity improvements of several orders of magnitude [86,87], re-examining these areal boundaries using next-generation, high-sensitivity imaging platforms will be important. Such advancements may further reveal fine-grained metabolic sub-networks that extend beyond the principal superior–inferior axis characterized in the present study.

Second, while the TACs of fPET-FDG are closely related to glucodynamics, their temporal variations do not directly reflect rapid glucose phosphorylation but rather a composite process involving transport across the blood-brain barrier, tissue uptake, and phosphorylation. Consequently, the extent to which observed fPET-FDG signal fluctuations capture true functional changes in tissue glucose uptake — particularly at sub-minute timescales — remains an open question, as these timescales deviate significantly from the steady-state assumptions that underlie FDG-PET metabolic mapping (i.e., when plasma FDG concentrations are equilibrated). Although our results indicate that the identified network patterns are dominated by slow, minute-scale variations, faster fPET dynamics still contribute to the overall network structure, and recent studies have reported robust oscillations in fPET TACs that appear to convey meaningful functional information with frame resolutions of a few seconds [54,88]. These faster dynamics could be more associated with glucose supply than to phosphorylation associated with glucose consumption [89] or to changes in blood flow. Disentangling these different contributions to glucodynamics will be important for accurate interpretation of network patterns characterized with fPET-FDG.

Finally, our study was conducted in a relatively young cohort, with a mean participant age of 19 years. As a result, the metabolic network organization characterized here may preferentially reflect features of late adolescent or early adult brain metabolism and may not fully generalize across the lifespan. Given the well-established age-related changes in cerebral metabolism and functional organization [50,90–92], extending these analyses to more diverse age groups will be important for assessing the developmental and aging trajectories of metabolic network architecture. In addition, the current sample size was modest (𝑁 = 24). While sufficient to reveal robust large-scale metabolic network patterns, future studies leveraging larger, openly available rsPET-MR datasets will be critical for refining the characterization of metabolic organization, improving statistical power, and enabling the identification of more subtle or heterogeneous metabolic sub-networks.

## Conclusion

In summary, we provide a detailed spatiotemporal characterization of the organizational patterns of resting-state metabolic connectivity using constant-infusion fPET-FDG. At local scale, connectivity-based boundary mapping identified abrupt transitions in metabolic connectivity profiles, shaped by both slow and fast fPET-FDG dynamics. At the global level, metabolic connectivity exhibits a robust superior-inferior axis that is consistently expressed in both network- and gradient-based representations and is primarily shaped by slow, minute-scale fPET-FDG dynamics. While these fPET-FDG-derived metabolic profiles diverge from traditional hemodynamic counterparts, they show clear alignment with known anatomical and energetic constraints. Together, these results demonstrate that fPET-FDG captures large-scale, metabolically grounded principles of cortical organization that are not accessible through BOLD-fMRI alone, thereby providing a complementary, metabolism-informed perspective for understanding intrinsic brain organization.

## Data and code availability

The dataset utilized in this study, Monash rsPET–MR is publicly available in Jamadar *et al.* [52]. The code used for preprocessing, analysis, and visualization is available at https://github.com/PenghuiDu/metabolic_connectivity.

## Author contributions

P.D. and J.E.C. conceived and designed the study. P.D. performed the formal analyses with support from S.E.C., T.X., and B.W.. J.E.C. and Q.L. funded and supervised this study. P.D. wrote the manuscript, all authors contributed to editing the manuscript.

## Acknowledgements

We would like to thank Dr. Alain Dagher, Dr. Bratislav Misic, and Mr. Moohebat Pourmajidian for sharing the genetic maps of energy metabolism pathways. This work was supported in part by the NIH (grants K99/R00-NS118120 and R21-MH135201), by the Harvard Mind Brain Behavior Faculty Research Award, by the Brain & Behavior Research Foundation Young Investigator Grant, by the BrightFocus Foundation Research Grant, by the Rappaport Foundation, and by the Australian Research Council Future Fellowship FT 250100206.

## Ethics declarations

### Conflict of interest

The authors declare no competing financial or non-financial interests that could have influenced the outcomes of this research.

### Ethical approval

The public human PET-MRI data analyzed in this study were acquired in accordance with the Australian National Statement of Ethical Conduct in Human Research. The original study that generated this dataset was reviewed and approved by the Monash University Human Research Ethics Committee, and all individuals provided informed consent.

## Notes

### Competing Interest Statement

The authors have declared no competing interest.

