## Supplementary Materials for "Human Cerebral Cortex Organization Characterized by Functional PET-FDG “Metabolic Connectivity”"

### Supplementary Methods

#### Boundary Mapping Refinement for Low-SNR fPET-FDG Data

To reduce artifacts caused by volume-to-surface resampling and smoothing on unevenly sampling surface meshes, a gradient regression based correction was developed and applied to both BOLD-fMRI and fPET-FDG boundary mapping, as illustrated in **Error! Reference source not found.a**.

*Surrogate Data Generation:* For each subject, surrogate datasets were created by random phase-shuffling of voxelwise time series, preserving temporal spectral properties while removing spatially structured connectivity. Both original and surrogate datasets underwent identical preprocessing steps, including volume-to-surface resampling and surface-based smoothing provided by Freesurfer [1] and Connectome Workbench [2].

*Connectivity and Gradient Estimation:* Vertexwise functional and metabolic connectivity were estimated using Pearson correlations followed by Fisher's Z-transformation on individual surface space. Local connectivity gradients were computed using the *cifti-gradient* function in Connectome Workbench [2] , following the approach of Gordon et al (2016) [3]. These gradients capture transitions in connectivity similarity across the cortical surface and form the basis for boundary detection.

*Gradient Regression Correction:* To account for artifactual effects, local gradients derived from the surrogate data were linearly regressed out from those of the original data on individual surface space. This step reduced smoothing and resampling-related biases while preserving genuine connectivity transitions.

*Boundary Detection:* Following correction, the local gradients were resampled onto the fsLR 32k [4] surface for group-level analysis. A 6mm FWHM surface smoothing was applied to the group-averaged gradients to improve SNR. The watershed-by-flooding algorithm [3,5] was then applied to the local gradient profile of each vertex to delineate connectivity boundaries and generate boundary probability maps. Adjacent parcels with weak shared boundaries were merged, vertices with high boundary uncertainty were masked, and small parcels (<15 vertices, ~30mm$^2$) were excluded to obtain the final surface parcellation.

#### Validation with Synthetic Data

The refinement for local gradient based boundary mapping was evaluated using synthetic datasets generated for each subject, from a spatially structured connectivity prior based on the Schaefer 7-network 100-ROI atlas [6], with ten times fPET data frames of each subject. A vertex-to-vertex connectivity matrix was constructed with correlations of 0.6 within parcels, 0.3 between parcels in the same network, and 0 between networks. Colored timeseries were simulated via Cholesky decomposition of the target correlation matrix and contaminated with Gaussian noise of varying amplitudes to emulate different SNR conditions (**Error! Reference source not found.b**).

Synthetic datasets were projected to the cortical surface and processed in the same way with the real data. As illustrated in **Error! Reference source not found.b**, without correction, spurious boundaries are produced, particularly in regions of high curvature (e.g., central sulcus, cingulate gyrus, superior temporal sulcus, and calcarine fissure) under low-SNR conditions. Gradient regression modestly reduced these artifacts and improved boundary precision under low-SNR conditions. When applied to pure Gaussian noise, the procedure did not generate structured boundaries, indicating that it did not introduce extra spurious patterns. Quantitative analyses (**Figure S4**) further confirmed that regression correction improved boundary detection at low SNR, though recovery remained limited relative to high-SNR data.


### Supplementary Figures

See next page.

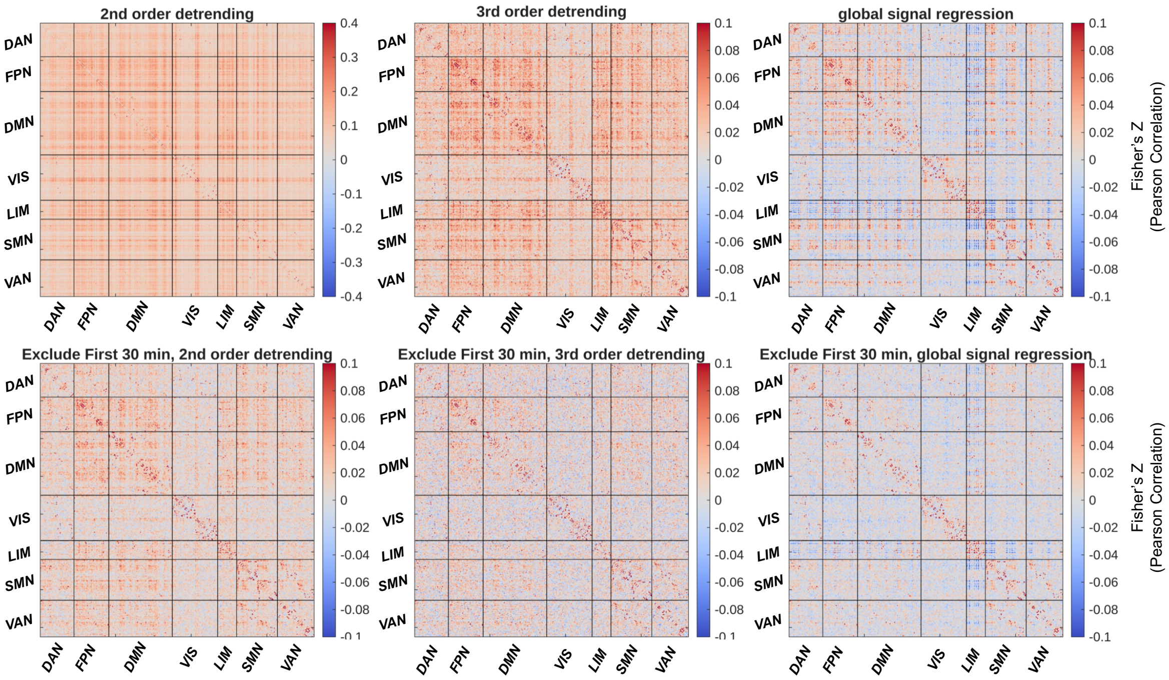


**Figure S1** Group-averaged MC matrices computed from fPET-FDG data using the Glasser 360-ROI parcellation, under different detrending and data inclusion strategies. The top row shows results including the whole session; while the bottom row excludes the first 30 minutes of scanning, during which tracer uptake has not likely reached pseudo-equilibrium. Each column corresponds to a different preprocessing approach: 2nd-order polynomial detrending, 3rd-order polynomial detrending, and global signal regression. Including the early time window introduces structured spurious patterns across all methods. Among the detrending strategies, 3rd-order detrending yields MC matrices with relatively fewer apparent structured patterns, while 2nd-order detrending leaves more prominent residual structure. Global signal regression introduces widespread anti-correlations in both cases. All connectivity values represent Fisher's Z-transformed Pearson correlations.

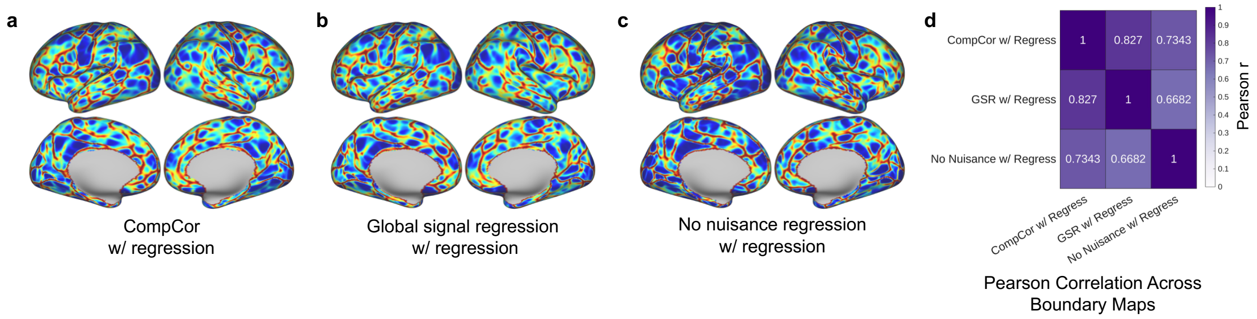


**Figure S2** Comparison of boundary mapping results across different fMRI processing strategies. (a-c) Boundary maps derived from different fMRI nuisance regression pipelines: (a) aCompCor, (b) global signal regression, and (c) no nuisance regression. Gradient regression was applied for all three pipelines before boundary mapping. (d) Spatial correlation matrix between boundary maps generated with different preprocessing approaches.

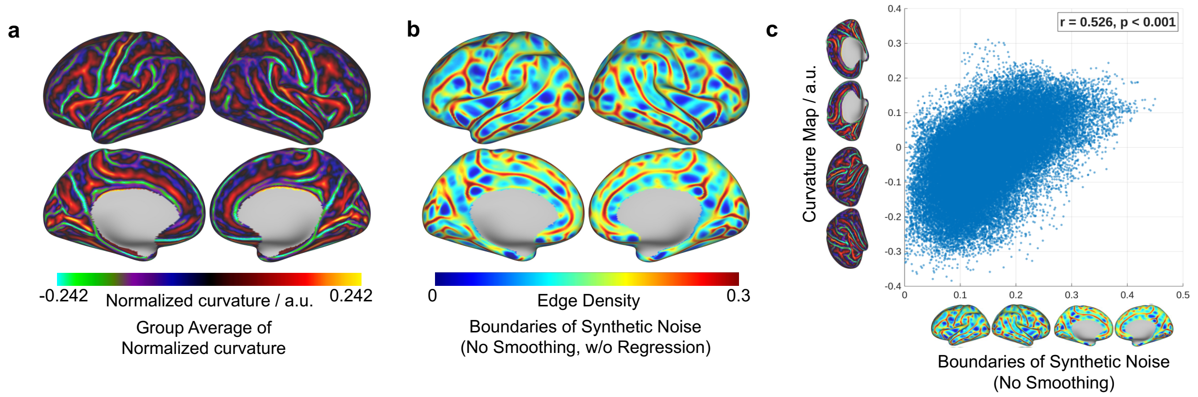


**Figure S3** Curvature-related artifacts in boundary mapping from synthetic noise. (a) Boundary maps derived from synthetic noise exhibit strong and spatially structured patterns, despite the absence of any true signal. These artifacts notably overlap with (b) regions of high cortical curvature. (c) Voxelwise correlation between curvature and the resulting boundary maps further confirms this effect, with significant, moderate to high positive correlations across the cortex ($r = 0.526,\ p < 0.001$), indicating that curvature-related confounds can induce spurious boundary structures even in the absence of true functional divisions.

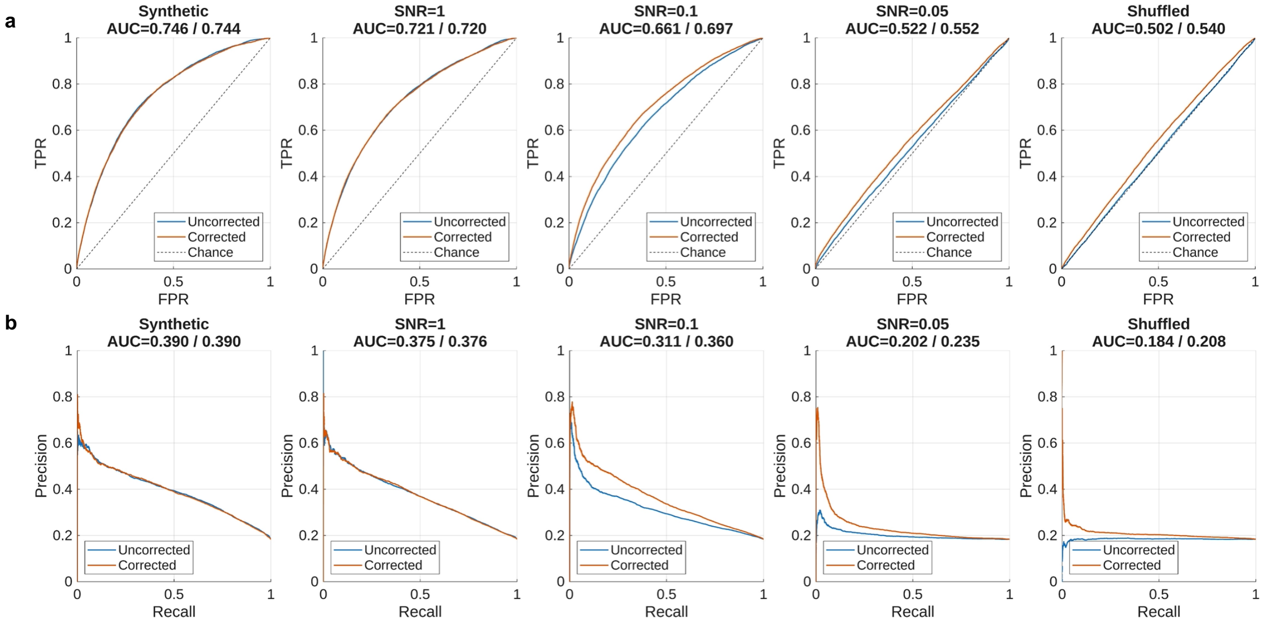


**Figure S4** Comparison of boundary mapping performance on the synthetic dataset with and without gradient regression. **(a)** ROC curves and **(b)** precision-recall curves are shown for the synthetic dataset under different SNR conditions, including a sham control with temporal phase shuffled. Each panel compares uncorrected (blue) and corrected (orange) models. AUC values are reported in each subplot (uncorrected vs. corrected with gradient regression).

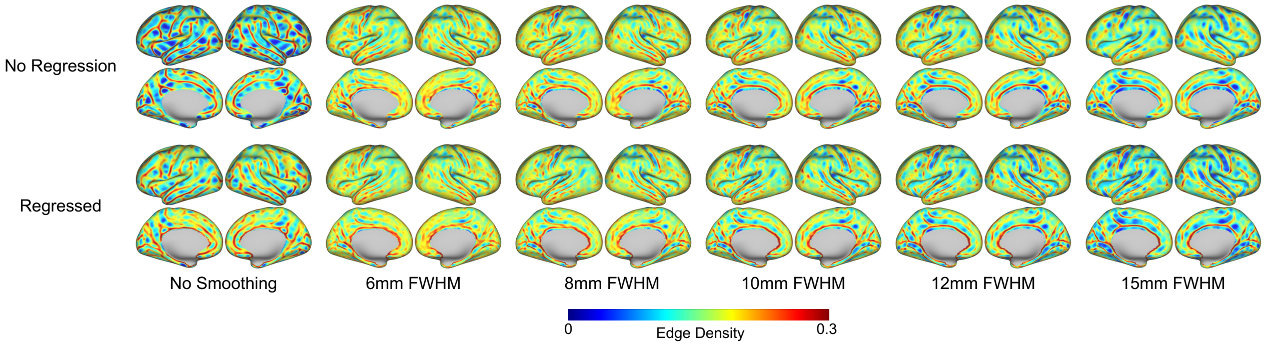


**Figure S5** Boundary maps computed from fPET-FDG data under varying spatial smoothing levels, with and without gradient regression. Each column corresponds to a different smoothing kernel size (left to right: no smoothing to progressively stronger smoothing). The top row shows boundary maps without gradient regression, and the bottom row shows the corresponding maps after applying gradient regression. Across smoothing levels, the combination of spatial smoothing and gradient regression progressively reduces structured artifacts related to cortical curvature, while preserving overall consistency in boundary patterns.

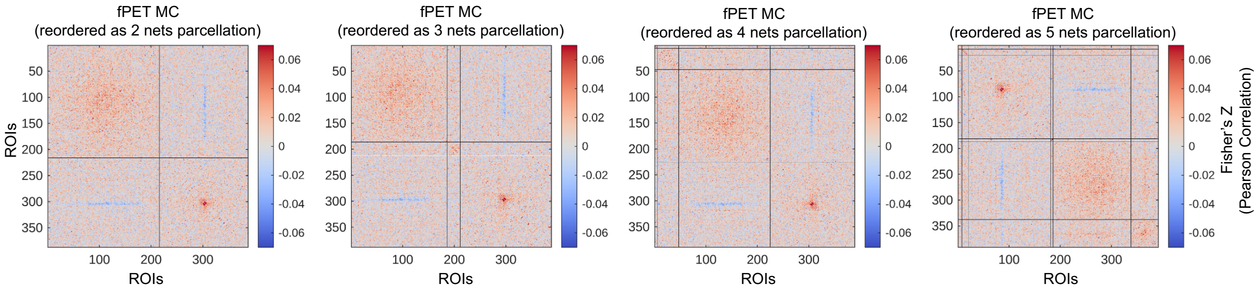


**Figure S6** Comparison of fPET-derived metabolic gradients across parcellation schemes and network-wise structure of the metabolic connectivity matrix. MC matrices computed from the study-specific parcellation are shown, reordered according to different numbers of detected functional networks. Each panel visualizes the same MC matrix with different reordering schemes.

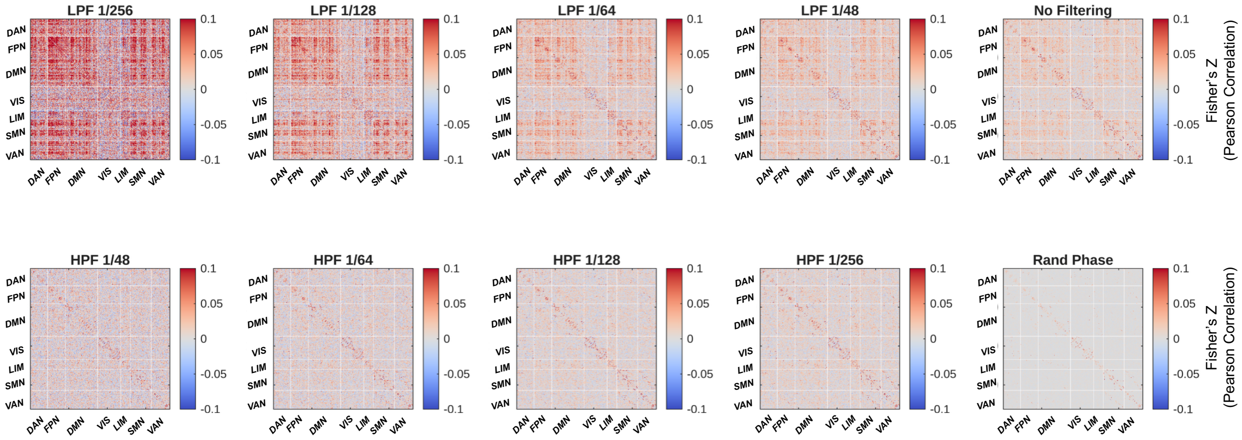


**Figure S7** Group-averaged MC matrices computed under different temporal filtering conditions, using the Glasser 360-ROI parcellation. The top row shows low-pass filtered data with progressively higher cutoff frequencies (from 1/256 Hz to 1/48 Hz), followed by unfiltered data ("No Filtering"). The bottom row shows high-pass filtered data with progressively lower cutoff frequencies (from 1/48 Hz to 1/256 Hz), followed by a random phase-shuffled control ("Rand Phase"). It is evident that MC patterns are strongly dominated by low-frequency components, with prominent structure retained even under very low-frequency cutoffs (e.g., 1/256 Hz, corresponding to ~4.3 minutes).

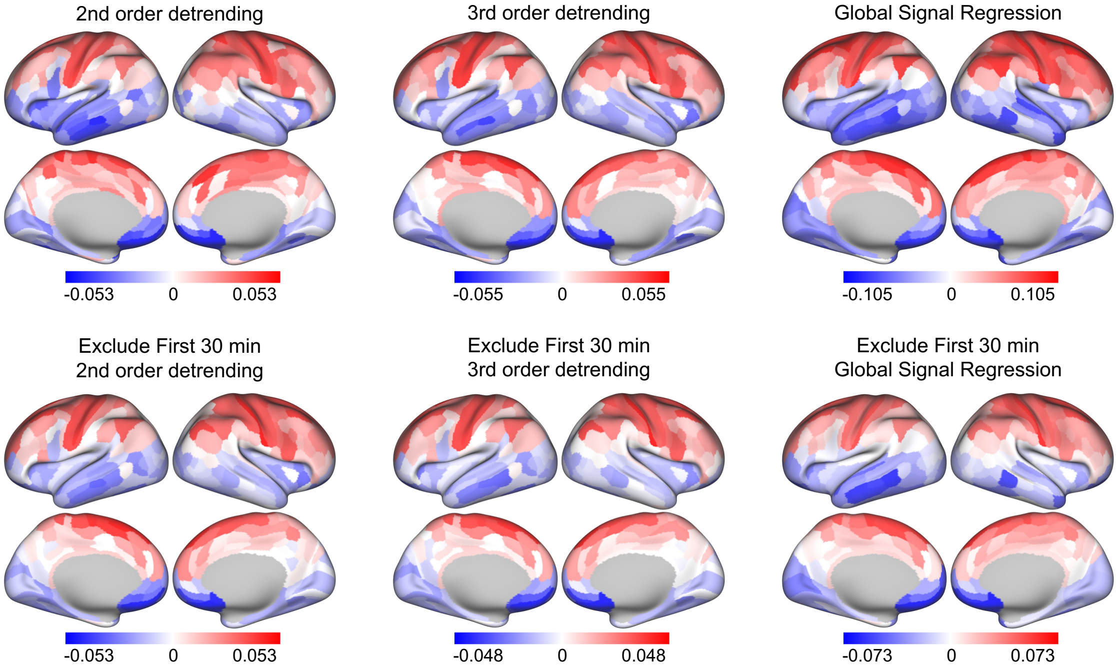


**Figure S8** First principal gradients computed from fPET-FDG metabolic connectivity under different detrending and data inclusion strategies. Similar to Figure S1, each panel corresponds to a specific detrending and data inclusioncondition, combining either 2nd-order detrending, 3rd-order detrending, or global signal regression, with or without excluding the first 30 minutes of data acquisition. Across all preprocessing pipelines, the resulting first gradient consistently exhibits a clear and spatially coherent superior-inferior axis, demonstrating the robustness of this macroscale organizational feature to processing variation.

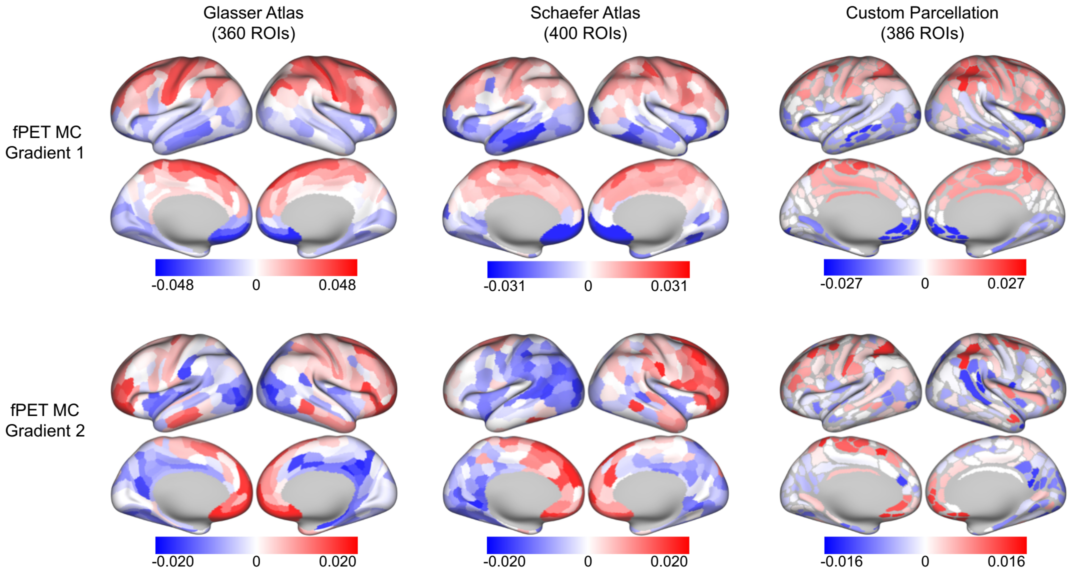


**Figure S9.** Principal gradients computed from fPET MC matrices under three different parcellation schemes: the Glasser 360-ROI atlas, the Schaefer 400-ROI atlas, and the study-specific parcellation (386 ROIs) in this work. General spatial patterns such as superior-inferior differentiation in gradient 1 are consistent across three schemes, but notable differences present particularly in gradient 2.
